# Coronary vessel assembly involves patterned endocardial sprouting and tip-cell-to artery specification

**DOI:** 10.1101/2022.12.20.517740

**Authors:** Elena Cano, Jennifer Paech, Masatoshi Kanda, Eric L. Lindberg, Irene Hollfinger, Caroline Brauening, Cornelius Fischer, Norbert Hübner, Holger Gerhardt

## Abstract

The coronary vasculature comprises superficial coronary veins and deeper coronary arteries and capillaries that critically support the high metabolic activity of the beating heart. Understanding how different endothelial progenitor sources jointly shape and remodel the coronary vasculature into its adult pattern has recently attracted great research interest, and spurred much debate. Here, using lineage tracing tools in combination with three-dimensional imaging, live-imaging in explants and single-cell transcriptional profiling, we demonstrate that sprouting angiogenesis drives both sinus venosus and endocardial contribution to coronary plexus. Whereas previous studies limit endocardial contribution to coronary vessels to the interventricular septum and ventral aspect of the heart, our study demonstrates extensive endocardial sprouting angiogenesis in the free ventricular walls. In particular, we identify a distinct subpopulation of endocardial cells that express future coronary markers and reside in both the embryonic and adult ventricular wall endocardium. Most importantly, we provide evidence for sprouting angiogenesis from both endocardium and subepicardial plexus towards the inner myocardial wall to determine pre-arterial specification. Additionally, sprouting from the endocardium leads to the establishment of perfused connections to the advancing coronary plexus, also followed by transitioning to the pre-arterial cell state. Distinct molecular profiles characterize sprouting populations in the intramyocardial and subepicardial layers that shape the prospective coronary arteries and veins, respectively. Harnessing the endocardial progenitors and targeting the distinct sprouting populations may in the future serve to tailor cardiac vascular adaptations for therapeutic purposes.

## Introduction

The coronary vasculature is the dedicated vascular system that supplies the heart. Despite its increasing clinical relevance, given the worldwide accelerated incidence of coronary artery disease (Timmis et al., 2022), the cellular mechanisms of coronary vasculature formation were not resolved until the last decade (Red-Horse et al., 2010, Wu et al., 2012). Since then, many studies have revealed intriguing particularities in the formation of the coronary vasculature. The coronary plexus develops as a mosaic of different embryonic sources that contribute simultaneously to the coronary endothelium (Cano et al., 2016; Carmona et al., 2020; Chen et al., 2014a; Katz et al., 2012; Red-Horse et al., 2010; Sharma et al., 2017; Tian et al., 2013a; Wu et al., 2012; Zhang et al., 2016), as well as to the mural cells that coalesce to complement the vessel wall (Cai et al., 2008; Chen et al., 2016; Volz et al., 2015; Zhou et al., 2008). Sinus venosus (SV), the inflow tract of the embryonic heart, and the endocardium, the inner endothelial layer lining the cardiac chambers, are the two predominant sources of coronary endothelium. SV-derived and endocardial-derived coronary vessels are believed to colonize mutually excluding regions of the heart (Chen et al., 2014a; D’Amato et al., 2022; Phansalkar et al., 2021; Sharma et al., 2017; Tian et al., 2013a). Interestingly, these two sources of coronary endothelial cells compensate each other when one of the sources is compromised (Sharma et al., 2017), suggesting a certain plasticity and redundancy of the progenitor pools, thus providing robustness to ensure the vascularization of the cardiac muscle. Nonetheless, the mechanisms that drive the SV-derived and endocardial-derived vessels to integrate and form a complete vasculature are not yet fully understood.

Several studies proposed endocardial angiogenesis as the mechanism driving endocardial contribution (Chen et al., 2014a; D’Amato et al., 2022; Sharma et al., 2017; Wu et al., 2012), however, lacking direct evidence. To gain deeper insight into potential links between origin, mechanisms of contribution and functional patterning, we have studied the endocardial contribution with high spatial resolution, combining various lineage tracing tools with three-dimensional and live imaging. We provide evidence for endocardial sprouting angiogenesis to be the process driving vascularization of the interventricular septum (IVS) as well as the ventral aspect of the heart. But we identify a wave of angiogenic endocardial activation that correlates spatiotemporally with the progression of the SV-derived plexus leading front at the dorsal and lateral aspects of the heart. Endocardial angiogenesis results first, in approximately 20% of the coronary endothelial cells (CoECs) of the free ventricular walls (FVWs) being recruited from the endocardium, and second, the establishment of endocardium-coronary plexus connections. Interestingly, these connections are lumenized and perfused and undergo an arterialization process. Using single-cell transcriptomics, we unraveled the transitory transcriptional states of the cells undergoing arterialization. This, together with the histological spatiotemporal mapping of the candidate pre-arterial markers, led us to propose that arterialization of the endocardium-coronary connections occurs through pre-arterial specification at the sprouting front during endocardial angiogenesis. Contrary to previous concepts on cardiac vascularization, our findings suggest that the heart uses the same common mechanisms already described for other vascular beds and organisms (Hasan et al., 2017; Hou et al., 2022; Luo et al., 2021; Pitulescu et al., 2017; Xu et al., 2014). We thus propose that prospective coronary arteries are also specified by this tip-cell-to-artery directed mechanism, since both SV-derived and endocardial-derived cells sprout to invade the inner segments of the myocardial wall.

## Results

### Coronary vasculature is initiated by angiogenesis at different sites of the developing heart

Our search for genetic tools to label coronary lineages revealed *Pdgfb* expression by the endothelium of the forming coronary plexus (fig 1A). We combined the use of inducible *PdgfbCreERT* mouse transgenic strain (Claxton et al., 2008) with the reporter *R26mTmG* strain (Muzumdar et al., 2007). We induced tamoxifen-mediated reporter labeling 24 hours before every timepoint analyzed, in order to genetically and irreversibly label by GFP the *Pdgfb*-expressing endothelial cells of the nascent coronary plexus (CoECs) (fig. S1A). To corroborate the CoECs identity of the *Pdgfb*-expressing cells, we confirmed the co-expression of Fabp4, a well-known marker of the coronary vasculature (Elmasri et al., 2009) (fig. S2C-D). At embryonic day (E)11.5, *Pdgfb* expression revealed a tip-cell specific pattern (Lindblom et al., 2003) (fig.1B), whereas from E12.5 onwards, it is upregulated by most CoECs (fig. 1A, fig. S2A-B). Minor fractions of endocardial cells (EndoECs) of the valves (fig. S2E) and ECs of the outflow tract (fig. S2F) also showed *Pdgfb* expression.

**Figure 1.**
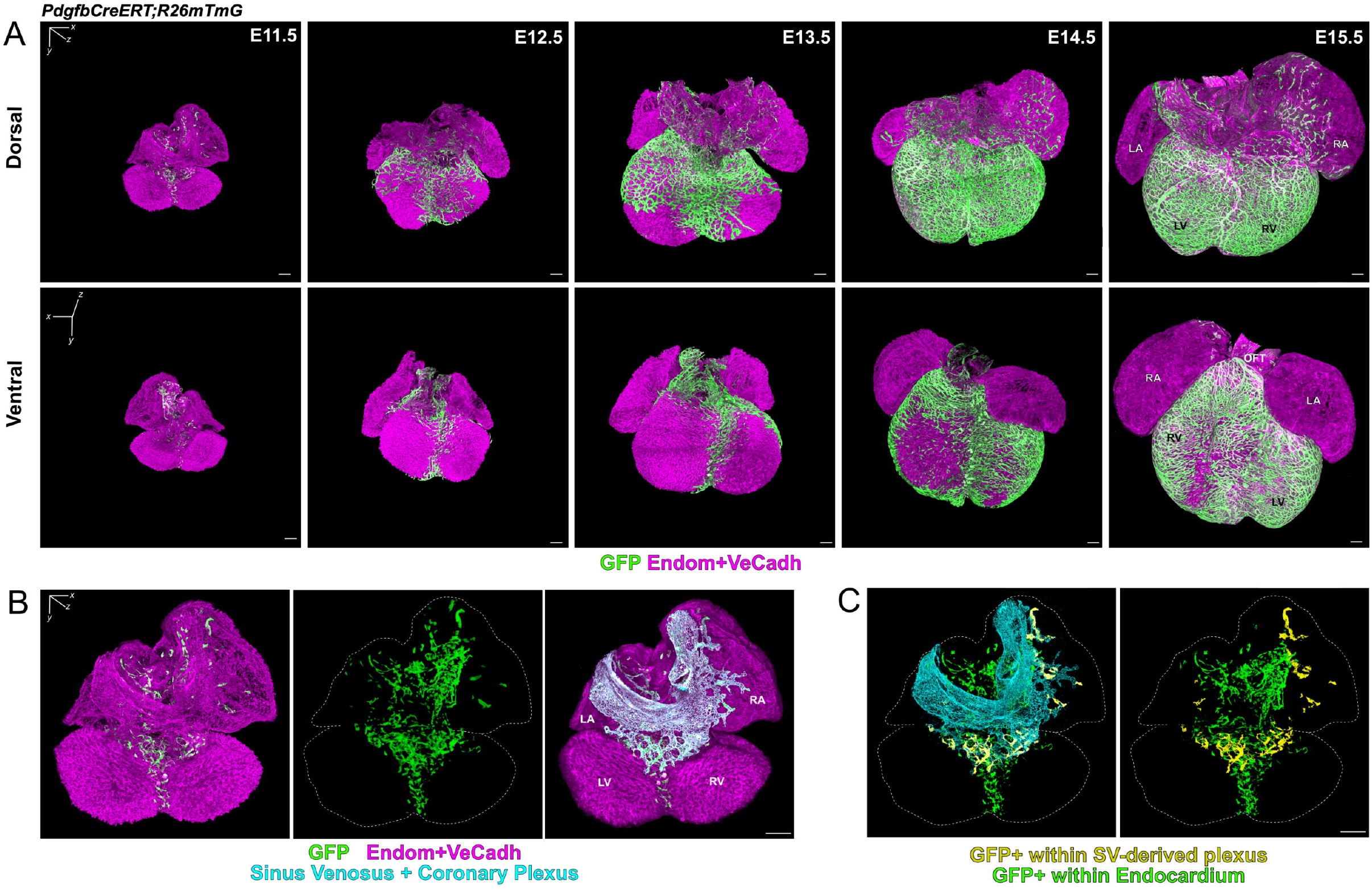
*Pdgfb*-expressing cells are found within coronary plexus and endocardium. A. Three-dimensional rendering of E11.5-E15.5 *PdgfbCreERT;R26mTmG hearts*. CoECs are labeled by GFP (green) while all cardiac ECs (CoECs and endocardial) are labeled with a cocktail of anti-VeCadherin (VeCadh) and anti-Endomucin (Endom) antibodies (magenta). Please note that GFP-expressing cells are also expressing VeCadh and Endom. B. The SV and the contiguous primary vascular plexus are highlighted in cyan by image post-processing segmentation. C. Not all *Pdgfb*-expressing ECs are included in the SV-derived plexus (yellow) but a significant number are within the endocardium (green). RA=Right Atrium; LA=Left Atrium; RV= Right Ventricle; LV=Left Ventricle; SV= Sinus Venosus; OFT= Outflow Tract. Scale bars=100μm.

Our three-dimensional imaging approach (fig. S1B) enabled us to identify the anatomical origin of nascent vessels, as well as the mechanistic processes driving their expansion. We identified two different sites where coronary vascularization is initiated.

First, at the dorsal side of the heart, an incipient plexus appeared at E11.5 within the subepicardial space of the atrioventricular canal (AVC) (fig. 1A, top row). This plexus extended from the SV, confirming angiogenic sprouting from the SV as the main endothelial source of the dorsal coronary plexus (fig. 1B) (Chen et al., 2014a; Red-Horse et al., 2010). The sprouting front expanded radially along the subepicardium to vascularize the ventricles, advancing the plexus behind this front to progressively vascularize the underlying compact myocardial layer.

Second, at the IVS, *Pdgfb*-expressing EndoECs invaded the forming septal myocardium and formed a local plexus (fig. 1C, S2G). The IVS plexus promptly anastomosed with SV-derived plexus as this one reached the IVS around E12.0 (fig. 1A). Additionally, the endocardium of the IVS also vascularized the ventral side of the heart (fig. 1A, bottom row). *Pdgfb*-expressing EndoECs sprouted outwards, invading either the myocardium or the subepicardial space, where they particularly formed endocardial buds (fig. S2H). These sprouts gained complexity, branching and establishing connections to finally form a local plexus that expanded laterally (fig. S2I). The ventral plexus anastomosed with the SV-derived plexus at the ventral area of both ventricles, closing the coronary network completely by E15.5 (fig. 1A).

Morphological and topological features of these dorsal *Pdgfb*-expressing EndoECs signified active sprouting, including tip cell-like phenotypes with numerous filopodia, and their longitudinal axis perpendicular to the endocardial lining, protruding towards the subepicardial space (fig. 2A).

**Figure 2.**
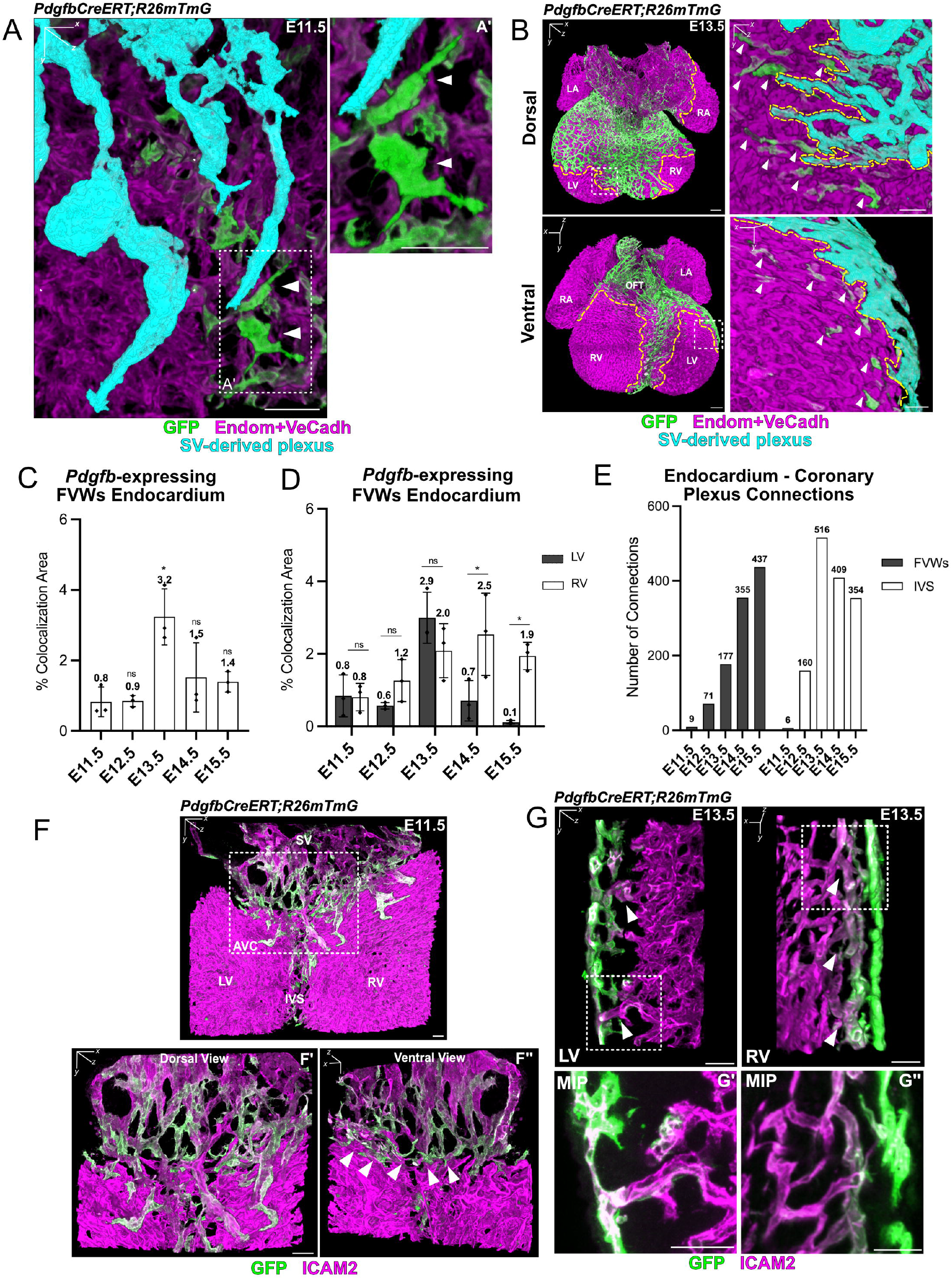
*Pdgfb*-expressing endocardial cells show signs of active angiogenic sprouting. A. Three-dimensional magnification of the sprouting front of forming SV-derived plexus (cyan) of a E11.5 *PdgfbCreERT;R26mTmG* heart. *Pdgfb*-expressing endocardial cells (arrowheads) show a tip cell-like phenotype (A’). B. Three-dimensional magnification of the sprouting front of forming SV-derived plexus (cyan) of a E13.5 *PdgfbCreERT;R26mTmG* heart. *Pdgfb*-expressing endocardial cells are located close to the sprouting front (arrowheads) in the FVWs. C. Percentage of endocardium that express *Pdgfb* in the FVWs along development. Quantified as GFP/Endom+VeCadh colocalization area (μm^2^), normalized to Endom+VeCadh total area, on the segmented endocardium of the FVWs (n=3). D. Percentage of endocardium (normalized GFP/Endom+VeCadh colocalization area) that express *Pdgfb* in RV vs. LV FVWs (n=3). E. Quantification of endocardial-coronary plexus lumenized connections in LV vs. RV along development (n=1). F. Three-dimensional magnification of SV-derived plexus of a E11.5 *PdgfbCreERT;R26mTmG* heart. Boxed area show the AVC from dorsal (F’) and ventral (F”) perspectives. Arrowheads point to ICAM2+ lumenized connections (magenta) between endocardium and incipient SV-derived plexus. G. Three-dimensional magnification of lumenized endocardial-coronary plexus connections (arrowheads) in RV and LV of a E13.5 *PdgfbCreERT;R26mTmG* heart. G1 and G2 are maximum intensity projections (MIP) of boxed areas. RA=Right Atrium; LA=Left Atrium; RV= Right Ventricle; LV=Left Ventricle; SV= Sinus Venosus; OFT= Outflow Tract, IVS=Interventricular Septum, AVC= Atriovenctirular Canal. Scale bars=100μm in B; 25μm in A, B boxed areas, F, G. Data are mean ± SD, ns=not significant, *=p<0.05, by Welch’s t-test in C; by Student’s paired t-test in D.

The three-dimensional mapping of the *Pdgfb*-lineage uncovered a remarkable mosaicism of the coronary vasculature, which is initiated at separate regions of the developing heart, yet with a similar angiogenic signature.

### *Pdgfb-expressing* EndoECs sprout and anastomose with the coronary plexus at the free ventricular walls

We have shown that, during the initial phases of coronary vasculature formation, *Pdgfb*-expressing EndoECs collectively respond in order to vascularize the IVS and the ventral aspect of the developing heat (fig. 1C). Interestingly, as development proceeds and the dorsal SV-derived plexus expands, we detected another contributing source of coronary endothelium: scattered *Pdgfb*-expressing EndoECs of the FVWs (fig. 1C, 2A-B).

The distribution of these *Pdgfb*-expressing EndoECs correlated spatiotemporally with the progression of the coronary plexus sprouting front (fig. 2B). Taking colocalization area (μm^2^) as an approximation of cell number, we estimated that a range of 1-3% of all EndoECs are upregulating *Pdgfb* in the FVWs at every timepoint studied, with a peak at E13.5 (fig. 2C). We also identified some regional differences in the distribution of these cells between left (LV) and right ventricles (RV), that indeed correlate with the vascularization process of the coronary plexus (fig. 2D). At E13.5, we observed a peak of *Pdgfb*-expressing EndoECs at the LV, which became rapidly vascularized over the next 24 hours. Similarly, at E14.5 and E15.5 we detected a higher presence of *Pdgfb-* expressing EndoECs in the RV, which vascularized slightly later than the LV (compare fig. 1A and fig. 2D). Together, these results unveil an angiogenic activation wave of the endocardium that correlated with the progression of the SV-derived plexus.

In general, sprouting vessels anastomose with either pre-existing vessels or adjacent sprouts to form a lumenized new branch or connection that expands the vascular network. Indeed, we also observed regular connections between the SV-derived coronary plexus and the endocardium. First, at E11.5, when the primary plexus appears at the AVC, SV sprouts and endocardial sprouts connected and formed a unique indistinguishable network (fig. 2F). Staining of the luminal endothelial membrane with ICAM2 demonstrated that these connections form a continuous lumen (fig. 2F). Later, as the SV-derived plexus expands, we also detected lumenized connections along both FVWs (fig. 2G). These lumenised connections between the ventricular chamber and the coronary plexus increased in number throughout development in the FVWs (fig. 2E). In the IVS, however, the number of endocardial-coronary plexus connections reached a peak at E13.5 and then decreased, suggesting that some of these connections may later be resolved (fig. 2E). This is consistent with the different timings of vascularization of the IVS versus the FVWs, as the endocardial-derived plexus of the IVS has been described to form before E13.5 (D’Amato et al., 2022). After the initial angiogenic phase, we observed some pruning of the established endocardial-coronary plexus connections (fig. 2E). In contrast, in the dorsal plexus, endocardial angiogenesis and EndoECs recruitment to the FVWs plexus occurred until the coronary vasculature was closed by E15.5, resulting in an increasing number of endocardium-coronary connections.

### Live imaging demonstrates sprouting and anastomosis of *Pdgfb-expressing* cells

In order to directly observe the dynamics of the angiogenic events that drive coronary morphogenesis, we performed live-imaging on *PdgfbCreERT;R26mTmG* E12.5 heart explants. *Pdgfb*-expressing cells emerged from the endocardium, as separate sprouts that initially did not form part of the SV-derived growing plexus. These acquired an exploratory and migratory tip-cell phenotype, showing multiple filopodia, and finally incorporated into the sprouting front (fig. 3).

**Figure 3.**
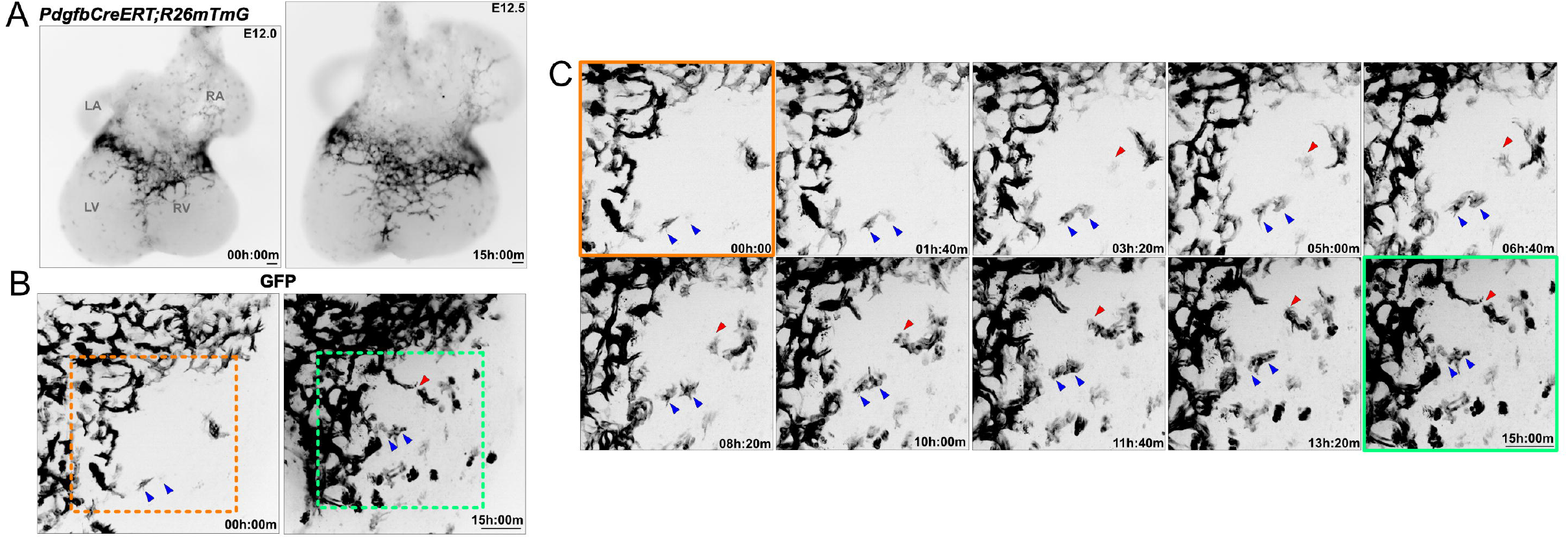
*Pdgfb-expressing* endocardial cells show exploratory and migratory phenotype and are actively recruited by the coronary plexus *ex vivo*. A. Dorsal view of a E12.0 *PdgfbCreERT;R26mTmG* explanted heart before (t=0) and after (t=15h) live-imaging. The dorsal coronary plexus (GFP, black) progression is comparable to the *in vivo* situation. B. Magnification of RV sprouting front before (orange boxed area) and after live-imaging (green boxed area). C. Time-lapse of SV-derived plexus sprouting front. *Pdgfb*-expressing endocardial cells (red and blue arrowheads) are actively attracted by the SV-derived plexus and finally make contact with its sprouting front. RA=Right Atrium; LA=Left Atrium; RV= Right Ventricle; LV=Left Ventricle. Scale bars=100μm.

### *Bmx*-lineage EndoECs significantly contribute to the coronary endothelium of the free ventricular walls

To confirm and quantify the contribution of the endocardium to the coronary vasculature, we used a second transgenic mouse line, *BmxCreERT;R26mTmG* (Ehling et al., 2013) as endocardial lineage tracing tool (fig. S1A). Our three-dimensional analysis of this line established that, when tamoxifen-mediated recombination is induced at E9.5, the *R26mTmG* reporter labels approximately 80% of all EndoECs at E12.5. In contrast, the SV is largely devoid of labelling; only up to 8% of all endothelial cells of the SV are labelled (fig. 4A-B), and are mostly found in the transition region where the SV drains into the right atrium (fig. 4A).

**Figure 4.**
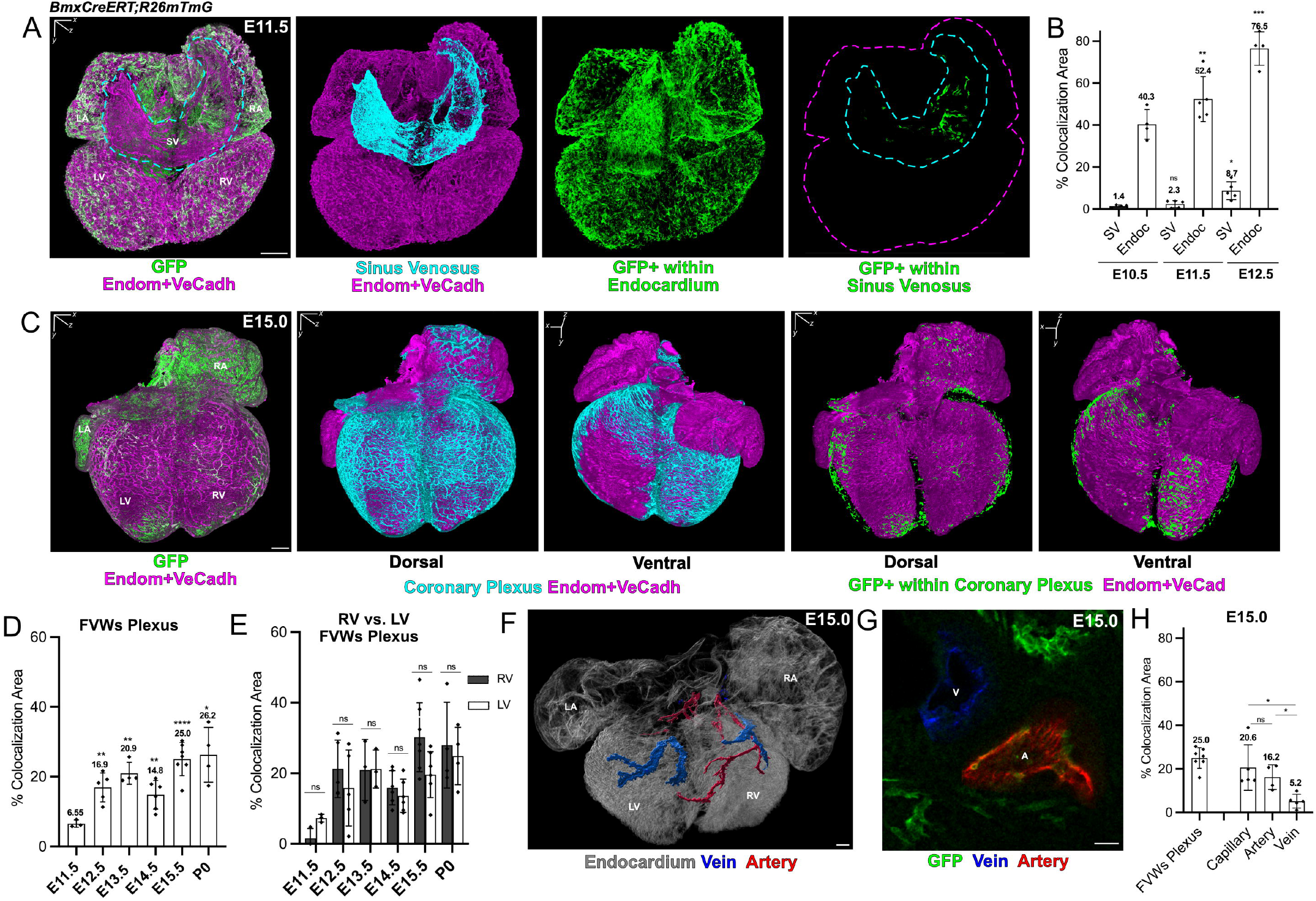
*Bmx*-lineage endocardial cells contribute differentially to coronary arteries and veins. A. Three-dimensional rendering of a E11.5 *BmxCreERT;R26mTmG* heart. SV is shown in cyan, while *Bmx*-lineage cells within the SV or endocardium are shown in green. *Bmx*-lineage cells found within the SV are located close to the transition area where SV drains to RA. B. Percentage of *Bmx*-lineage cells within SV or endocardium within the tamoxifen recombination window (E9.5-E12.5). Quantified as GFP/Endom+VeCadh colocalization area (μm^2^), normalized to Endom+VeCadh total area, on the segmented SV and endocardium (n=4-5, see dots on bars). C. Three-dimensional rendering of a E15.0 *BmxCreERT;R26mTmG* heart. *Bmx*-lineage derived CoECs (green) distribute broadly within the coronary plexus (cyan) of the ventricles. D. Percentage of coronary endothelium (normalized GFP/Endom+VeCadh colocalization area) of the FVWs plexus that derive from the *Bmx*-lineage (n=4-5, see dots on bars). E. Percentage of coronary endothelium (normalized GFP/Endom+VeCadh colocalization area) of the RV vs. LV FVWs that derive from the *Bmx*-lineage (n=3-7, see dots on bars). F. Post-processing segmentation of developing coronary arteries (red) and veins (blue) in a *BmxCreERT;R26mTmG* E15.5 heart. G. Single z plane of heart shown in F*. Bmx*-lineage cells (green) in coronary artery (red) and veins (blue). H. Percentage of coronary endothelium (normalized GFP/Endom+VeCadh colocalization area) that derive from *Bmx*-lineage in FVWs plexus, only capillaries, only arteries or only veins (n=4-7, see dots on bars). RA=Right Atrium; LA=Left Atrium; RV= Right Ventricle; LV=Left Ventricle; SV=Sinus venosus; Endoc=Endocardium, FVWs=Free ventricular walls; V= vein; A= artery. Scale bars=100μm in A, C, F; 25μm in G. Data are mean ± SD, ns=not significant, *=p<0.05, **=p<0.01, ***=p<0.001, ****=p<0.0001 by Welch’s t-test in C; by Student’s paired t-test in D.

Since Bmx is also known to be expressed by arterial cells (Ehling et al., 2013), we determined the time window in which a single tamoxifen injection leads to recombination. This is essential to be able to distinguish between bona fide endocardial lineage tracing, or potential later recombination events during differentiation of coronary arteries. First, we examined the subcellular localization of the CreERT recombinase by immunolocalization using an anti-estrogen receptor alpha (ERα) antibody. Second, since the *R26mTmG* cassette, upon recombination, excises the tdTomato (tdTom) coding sequence while triggering the transcription of the GFP coding sequence, we assessed the GFP fluorescence increase and the tdTom fluorescence decrease upon tamoxifen administration (fig. S3). During the first 48 hours post-tamoxifen (E10.5-E11.5), CreERT was localized exclusively in the nuclei of *Bmx*-expressing cells. Only within this time window, did we observe cells with low levels of GFP expression, an indication of a recent recombination event (fig. S3A-B). Most GFP-expressing cells were still showing tdTom fluorescence at this stage. From E12.5 onwards, CreERT was localized in the cytosol of the *Bmx*-expressing cells (fig. S3C-E). Some of these cells showed cytosolic CreERT but no GFP reporter activation (fig. S3C), indicating that recombination was no longer taking place. At E12.5, only a few of the GFP-expressing cells were still tdTom positive (fig. S3C). *Bmx*-lineage capillary CoECs of both early (fig. S3D) and mature plexus (fig. S3E), expressed GFP, but no tdTom nor CreERT, indicating they either were derived from (by division) – or directly represented (by sprouting) - endocardial *Bmx*-lineage cells (fig. S3D). On the other hand, arterial *Bmx*-lineage CoECs did express CreERT, although exclusively located in the cytosol, thus confirming that tamoxifen was no longer actively driving recombination at this time point (Chen et al., 2016) (fig S3E). These results validate the *BmxCreERT* driver as endocardial-specific when tamoxifen is administered at E9.5 (fig. S3F).

We further assessed whether the *Bmx*-expression by arterial CoECs autonomously triggers CreERT-mediated recombination by administration of tamoxifen at E13.5 (fig. S4A). Analysis at E15.5 revealed that the contribution of the *Bmx*-lineage endocardium to all coronary vasculature compartments, i.e. plexus, arteries and veins was reduced to 2-5% (fig. S4B). This indicates first, that *Bmx* upregulation by arterial cells occurs at later gestational stages, and second, that most of the endocardial contribution occurs before this developmental stage. Curiously, we detected endocardial-derived αSMA-pericytes and αSMA+ vascular smooth muscle cells (Chen et al., 2016) (fig. S4C). This contribution was never observed when tamoxifen is administered at E9.5, again providing evidence for tamoxifen to be cleared out of the circulation at least by E13.5.

After confirming the specificity and validity of our endocardial lineage tracing tool and methodology, we quantified the contribution of *Bmx*-lineage EndoECs to the coronary plexus. Such endocardial-derived CoECs accounted for ~20% of total CoECs of the FVWs (excluding IVS) at birth (fig. 4C-D) and were broadly distributed in LV and RV (fig. 4E).

However, when we analyzed the distribution of endocardial-derived CoECs within different coronary vessels type, i.e. arteries, veins and capillaries (fig. 4F), we observed a markedly differential contribution (fig. 4G-H). At E15.0, whereas microvascular and arterial vessels were comprised of 16-20% of endocardial-derived CoECs, veins showed only a minor contribution of just 5% (fig.4H). These data demonstrate a significant endocardial contribution to CoEC at the FVW, yet with a strong bias towards coronary arteries and capillaries, rather than coronary veins. In other words, the endocardium harbors a significant number of cells that function as progenitors for coronary vessels, in particular for coronary capillaries and arteries.

### Single cell-sequencing identifies coronary endothelial diversification throughout development

In order to identify the transcriptional identity of these endocardial progenitors of coronary vessels and to decipher whether endocardial-derived endothelial CoECs bear a distinctive endogenous transcriptional profile, we performed single-cell RNA sequencing on FACS-sorted cardiac endothelial fractions (CD31+/CD45-) from four different stages: E12.5, E15.5, postnatal day (P)2 and adult (8 weeks old) (fig. S1A). Rigorous quality control was performed and no sex batch effect was detected. Non-endothelial cells, immediate-early gene-expressing ECs and lymphatic ECs were removed from the analysis.

Unsupervised clustering using Uniform Manifold Approximation and Projection (UMAP) method on the combined datasets resulted in four big groups of clusters that we assigned as endocardium, coronary endothelium, proliferating cells and valvular endocardium, based on the expression of the top differential expressed genes. Thus, we identified five clusters of endocardial cells, referred to as Endoc I to Endoc V; five clusters of coronary endothelium, referred to as CoEC I to CoEC V; three proliferating clusters, referred to as Prolif I to III, and finally, two endocardial clusters related to the valves, referred to as Valve I and Valve II (fig. 5A). All these clusters were annotated (fig. 5C-D) upon validation of the differential expressed genes using the *in situ* hybridization GenePaint database (Visel et al., 2004) (fig. S6). The distribution of all observed cell types recapitulated the expected composition of the cardiac tissue, where the endocardial fraction and proliferative endothelium were predominant during embryonic stages and the CoEC fraction was predominant in the postnatal stages (fig. 5C, second column). Also, while all endocardial clusters distribution appeared to be stable across all stages studied (fig. 5E), CoECs showed an increasing diversity from development to adulthood, corresponding to the specification of the different vessel types and their maturation state (fig. 5F).

**Figure 5.**
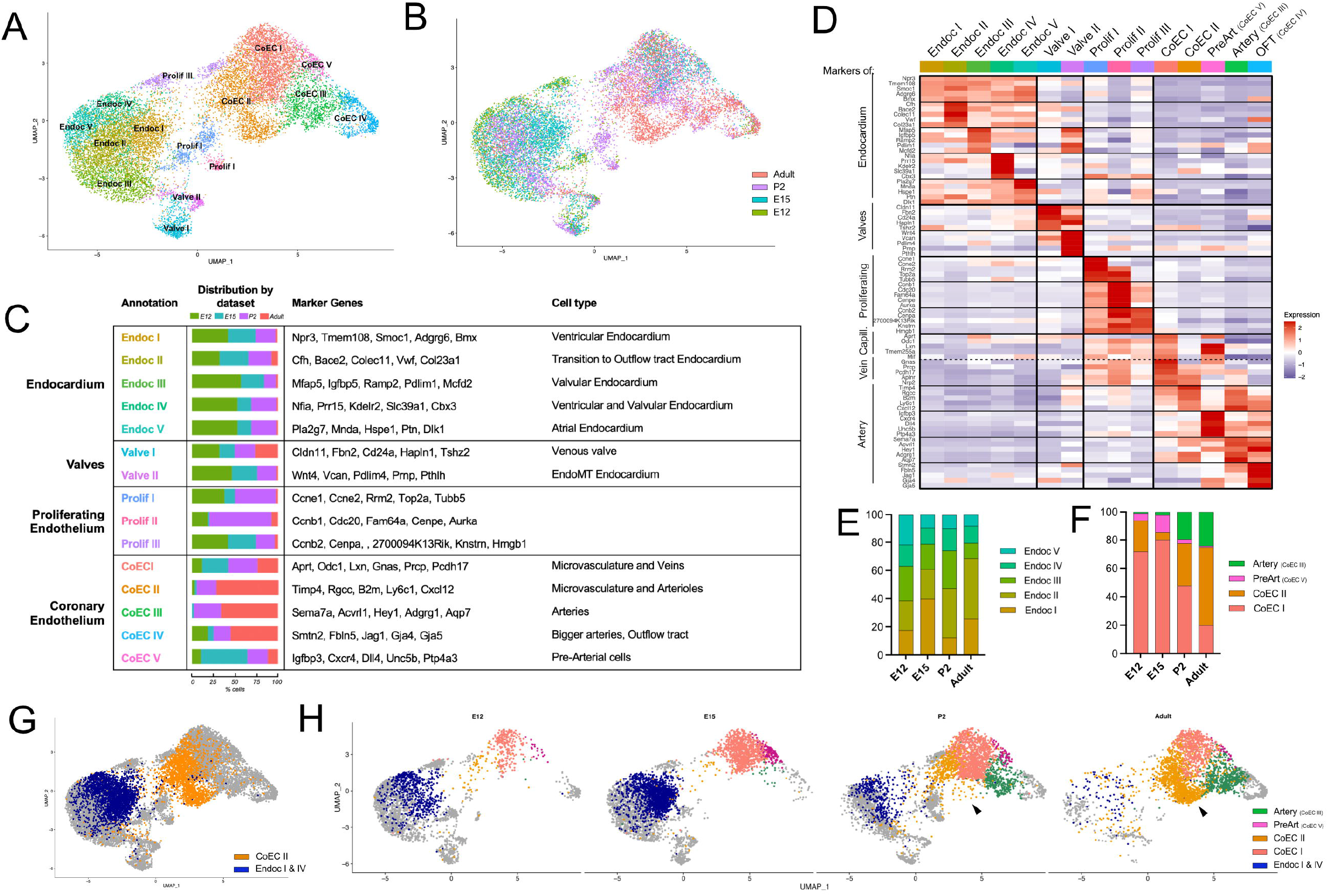
Combined E12-E15-P2-Adult scRNAseq identifies potential endocardial progenitors as well as increasing coronary endothelium diversity. A. UMAP plot of cardiac ECs from E12-E15-P2-Adult hearts combined datasets, grouped in 4 subpopulations, each composed by various clusters: coronary endothelium (CoEC I – CoEC V), endocardium (Endoc I-Endoc V), valvular endocardium (Valve I and II) and proliferating cells (Prolif I-III). The numerical order of the annotations correspond to the size in cell number of every cluster within each subpopulation. B. Previous UMAP plot color-coded by the dataset of origin (i.e. developmental stages). C. Annotations of every cluster (first row), with their distribution throughout the different developmental stages (second row), top marker genes (third row) and the assigned cell type after gene marker validation (last row and fig. S6). D. Heatmap of top marker genes in all annotated clusters. E. Distribution of all endocardium clusters in every studied developmental stage. F. Distribution of all coronary endothelium clusters in every studied developmental stage. G. UMAP plot where Endoc I and Endoc IV (blue) and CoEC II (orange) clusters were highlighted. H. UMAP plot split by developmental stage, where Endoc I and Endoc IV (blue) and all coronary endothelium clusters are highlighted: CoEC I (capillaries-to-veins, light pink), CoEC II (capillaries-to-arterioles, orange), CoEC III (arteries, green) and CoEC V (pre-arterial cells, dark pink). RA=Right Atrium; LA=Left Atrium; RV= Right Ventricle; LV=Left Ventricle.

At the initial phase of coronary vasculature formation (E12) (fig. 5F,H), the immature plexus was dominated by a microvascular subtype, the CoEC I (72% of all CoECs - without CoEC IV, which are outflow tract cells) which showed a capillary-to-venous transcriptional profile. We also detected some cells of CoEC II cluster, with a capillary-to-artery transcriptional profile (22%). In addition, we identified cells of cluster CoEC V (5%), that matched the recently identified pre-arterial cell profile (Hou et al., 2022; Luo et al., 2021; Su et al., 2018). We further observed cells of CoEC IV cluster, with a clear mature artery profile, which we assigned to cells of the outflow tract (OFT, aorta and pulmonary artery). This OFT cluster (CoEC IV) was stable throughout all stages studied in terms of its transcriptional profile, distribution and cell number (fig. 5H).

At E15 (fig. 5F,H), the coronary plexus showed signs of arterial remodeling, reflected by the appearance of first Arterial cells (CoEC III, 2%). Also, the Pre-arterial (CoEC V) cell number increased, reaching its peak at this stage (12%). At postnatal stages, both the capillary-to-veins CoEC I cluster (47%) and the Pre-artery cluster (CoEC V) decreased (3%), while the CoEC II (30%) and Artery (CoEC III) clusters (24%) increased (fig. 5A-D). At adult stages, the CoEC II (55%) cluster became predominant and also underwent further specification with the appearance of a new cluster subgroup (arrowheads in fig. 5D).

The histological validation of the candidate markers of every CoEC cluster (fig. S6H-M), suggested that CoEC I cells, which showed a capillary-to-vein profile, are cells located in the subepicardium at embryonic stages, as well as in the IVS. Further categorization of this cluster by analysis of already known markers (Phansalkar et al., 2021), served to identify IVS capillary, subepicardial capillary and vein subclusters within the CoEC I cluster (fig. S6H-J). On the other hand, the validation suggested that CoEC II cells map to an intramyocardial location (fig. S6K-L). At embryonic stages, cluster CoEC II expressed a capillary profile (fig. S6K). However, at postnatal stages, we detected a new branch of this cluster, which profoundly increased at the adult stage. This branch was defined by the expression of both capillary markers such as *Car4*, and arterial markers, such as *Cxcl12* (fig. S5A, S6M-N). Their location and validated transcriptional profile were consistent with intramyocardial capillary cells that underwent postnatal arterialization.

### Single cell-sequencing identifies putative endocardial progenitors of CoECs

Comparing all clusters across the developmental stages, we could not identify a cluster of CoEC that clearly retained any imprinted signature of the embryonic progenitor pool they were derived from (i.e. endocardium or SV). However, we observed cells mapping at intermediate positions (UMAP) between coronary and endocardial clusters. In particular, we observed overlap between EndoECs belonging to Endoc I and Endoc IV clusters, and the CoEC II cluster (fig. 5G, S5B-C). According to our annotation, Endoc I and Endoc IV clusters corresponded to EndoECs of the ventricular endocardium (fig. S6A,D), while CoEC II cells were found at the inner myocardial wall (fig. S6K-L) and show a capillary-to-artery transcriptional profile (fig. 5D). This pattern would be consistent with endocardial progenitors predominantly populating the inner myocardial wall (Chen et al., 2014a; Wu et al., 2012), contributing to capillaries and arteries rather than to veins (fig. 4H).

The GO-term analysis of Endoc I and Endoc IV revealed an enrichment of genes involved in cell migration (fig. S5D), in agreement with the actively exploratory phenotype observed by the endocardial-derived cells during live-imaging (fig. 3). The analysis of the differentially expressed genes of Endoc I and Endoc IV (fig. S5E), in comparison with the rest of endocardial clusters, identified *Fabp4*, a well-known marker of CoECs (Elmasri et al., 2009), as a distinct marker of these two clusters of ventricular EndoECs. In addition, both clusters showed higher expression of *Actb*, upregulation of which would signify a high actin turnover, likely due to higher cell motility. Staining and analysis on three-dimensional wholemounts confirmed the presence of scattered cells within the endocardium that distinctively expressed Fabp4 (fig. S5F). These data suggest that the endocardium harbors a distinct cluster of progenitors of CoECs, which can be identified by Fabp4 expression, and which become activated to engage in sprouting angiogenesis to shape an interconnected coronary vascular plexus. Interestingly, this population remained visible in the scRNAseq data also in the adult samples (fig. 5G-H).

### Pre-arterial cells share tip cell markers and persist in the adult

Pre-arterial cells have been defined as SV-derived venous cells that switch on an arterial specification program, abruptly upregulating mature arterial markers, such as Connexin40 (*Gja5*, Cx40), to subsequently build coronary arteries. Since coronary arterial remodelling is triggered by the onset of blood flow, and pre-arterial cells are detected prior to the connection of the coronary plexus to the aorta, which initiates perfusion (Chen et al., 2014b; Dyer et al., 2014; Tian et al., 2013b), pre-arterial cells have been assumed to signify pre-specified progenitors of arteries (Su et al., 2018; Trimm & Red-Horse, 2022).

Indeed, as early as E12, we identified that CoEC V/PreArtery cluster featured a typical capillary transcriptional profile, but also multiple arterial markers (fig. 6A). According to our data, and in contrast to what has been described for pre-arterial cells (Su et al., 2018), pre-arterial cells did not show significant expression of venous markers (fig. 6A). Curiously, these cells also showed a considerable enrichment of tip cell markers (fig. 6A).

**Figure 6.**
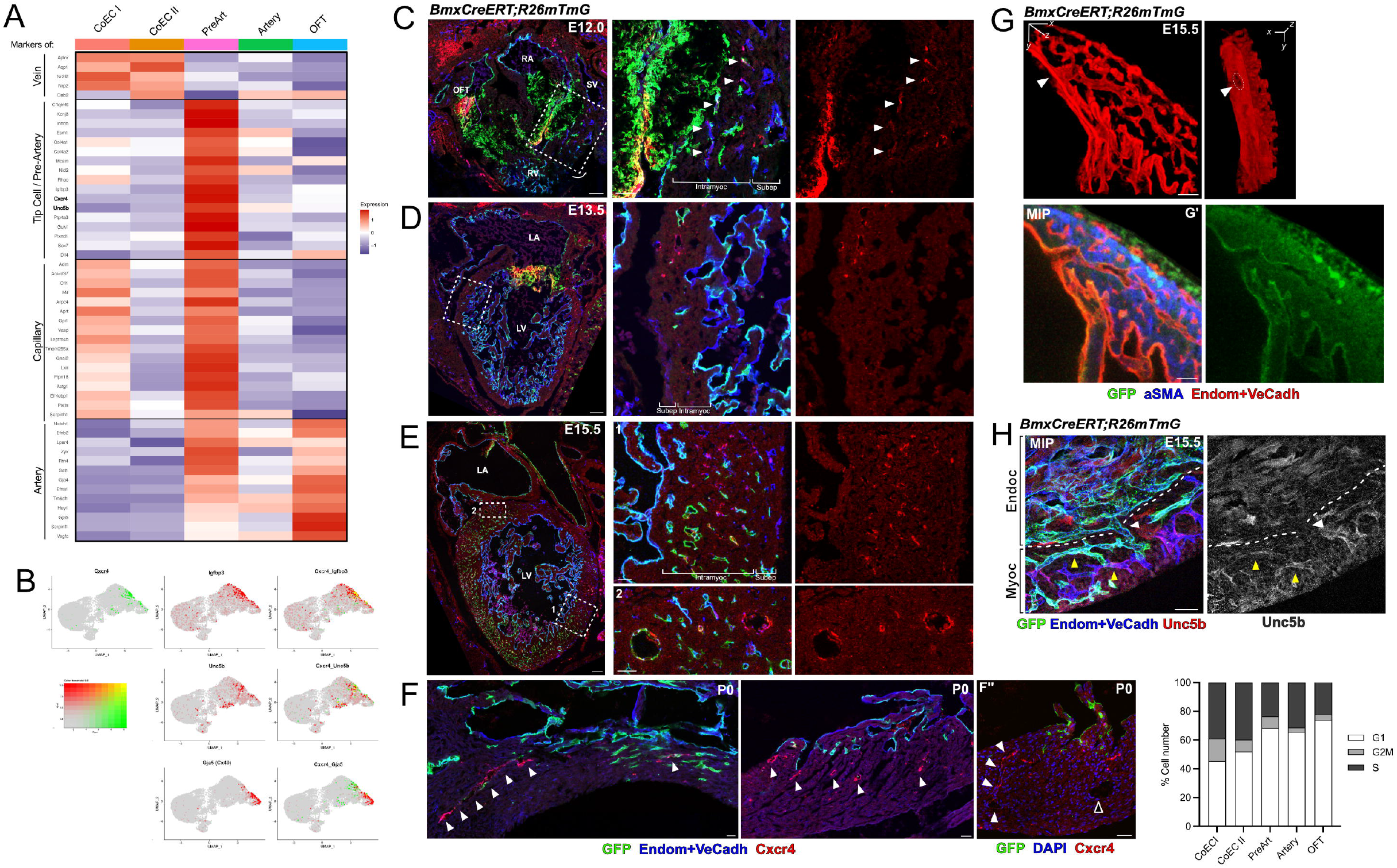
Endocardial-coronary plexus connections undergo arterialization. A. Heatmap of top marker genes in coronary endothelium clusters. B. Gene expression profile of *Cxcr4* versus the pre-arterial markers *Igfbp3* and *Unc5b* and the arterial marker *Gja5*. C-E. Sections of E12.0-E15.5 *BmxCreERT;R26mTmG* hearts stained against Cxcr4 (red) and Endom and VeCadh (blue). Arrowheads point to Cxcr4-expressing cells in the inner section of the myocardial wall. F. Sections of P0 *BmxCreERT;R26mTmG* hearts stained against Cxcr4 (red) and Endom and VeCadh (blue) or DAPI (blue, in F’). Arrowheads point to Cxcr4-expressing cells in the subendocardial area. Black arrowhead in G points a Cxcr4-mature artery. G. Three-dimensional rendering and maximum intensity projection (MIP, G’) of a segment of the FVW of a E15.0 *BmxCreERT;R26mTmG* heart stained against Endom+VeCadh (red) and aSMA (blue). Endocardium directly connects (arrowhead) with a coronary vessel located at the inner ventricular wall and that show an arterial phenotype. H. Maximum intensity projection of a segment of the FVW of a E15.0 *BmxCreERT;R26mTmG* heart stained against Unc5b (red) and Endom and VeCadh (blue). Endocardium directly connects (arrowhead) with coronary vessels located at the inner ventricular wall that show Unc5b expression. I. Quantification of the fraction of cells undergoing different cell cycle phases per cluster. RA=Right Atrium; LA=Left Atrium; RV= Right Ventricle; LV=Left Ventricle. SV=sinus venous. Scale bars=100μm in C-E; 50μm in F, G, H; 25μm in C boxed areas, G and H.

In order to further characterize their emergence and distribution, we performed spatiotemporal mapping of Cxcr4 and Unc5b immunolocalization, as they are specific markers of pre-arterial cells (fig. 6A-B). Since the early phases of coronary vasculature development, we found Cxcr4 to be only expressed by cells of the plexus that had already invaded the compact myocardial wall, when both subepicardial and intramyocardial plexus were well distinguished (fig. 6C-D). As the coronary plexus progressed, Cxcr4-expressing and Unc5b-expressing capillaries were found extensively within the intramyocardial segment of both ventricular walls (fig. 6E, fig. S7). In contrast, in those areas where remodeling arteries were observed, Cxcr4 and Unc5b expression became restricted to arterial cells and absent from the surrounding capillaries (fig. 6E, fig. S7A). This pattern would be consistent with pre-arterial cells of intramyocardial capillaries forming coronary arteries.

We also observed that mature Cx40-expressing artery cells lacked Cxcr4 expression (fig. 6B,F’’). Cxcr4 and Cx40 expression only overlapped in a minimal fraction of pre-arterial cells (fig. 6B), which we believe reflects pre-arterial cells transitioning into Cx40-expressing arteries. In contrast to what has been described previously (Raftrey et al., 2021; Su et al., 2018), we found that Cx40 was not expressed at early stages of plexus formation (E12) (fig. S5G).

Interestingly, in neonates, Cxcr4-expressing cells were found in the subendocardial zone (fig.6F). Often, these Cxcr4 and Unc5b-expressing cells were found in arterioles, which extend from the endocardial lumen (fig. 6G-H). Given the position of the Cxcr4-expressing cells, and the wider caliber, these connections between endocardium and coronary vessels appeared to undergo arterialization.

Indeed, pre-arterial cells were detected throughout embryonic, postnatal and adult stages exhibiting a quite stable transcriptional profile (fig. 5F, H). Interestingly, these data would suggest that pre-arterial cells do not become depleted from the heart even after the onset of blood flow and the definitive specification into mature arteries.

### Coronary plexus is perfused and remodeled before the connection to the aorta

The onset of perfusion of the coronary vasculature is widely accepted to occur when the growing coronary plexus connects to the aorta with the formation of the so-called coronary ostia, an event that triggers the onset of blood flow and subsequently network remodeling (Cavallero et al., 2015; Chen, et al., 2014b; Dyer et al., 2014; Tian et al., 2013b). Support for this concept comes from the observation that remodeling reportedly does not take place in mutants that fail to form the coronary ostia (Cavallero et al., 2005; Chen et al., 2014b; Gonzalez-Iriarte et al. 2003; Icardo et al. 2001; Ivins et al., 2015; Tomanek et al., 2008; Walker et al. 2005). Given that the appearance of pre-arterial cells occurs before the assumed perfusion of the coronary vasculature, their specification is believed to be uncoupled from blood flow (Su et al., 2018; Trimm & Red-Horse, 2022).

We studied the perfusion and the remodeling process of the coronary plexus in a three-dimensional manner and observed that the coronary plexus showed clear signs of perfusion since the beginning of its formation (fig. S8). Already at E11.5, ICAM2 staining demonstrated that the coronary plexus is continuously lumenized. Also, red blood cells could be readily observed within the dorsal plexus, as well as within the incipient endocardial buds that contributed to the ventral plexus (fig. S8A). At E12.5, the right coronary ostium was formed and lumenized, but no signs of remodeling were observed yet in the proximal coronary stem nor in the intramyocardial plexus. Remarkably, we already observed signs of remodeling in the subepicardial plexus, where the prospective right coronary vein could be identified (fig. S8B). Already at this stage, first pre-arterial cells were detected in the inner myocardial wall (fig. 5H, 6C) (Su et al., 2018). However, despite this early pre-arterial specification, remodeling of the coronary stems, as well as of the proximate intramyocardial plexus, was only detected after both coronary ostia were fully formed at E13.5 (fig. S8C).

## Discussion

The coronary vasculature develops as a mosaic of cells from different tissue sources (Cano et al., 2016; Carmona et al., 2020; Chen et al., 2014a; Chen et al., 2016; Katz et al., 2012; Red-Horse et al., 2010; Tian et al., 2013a; Wu et al., 2012). It is well accepted that endocardium and SV lineages contribute differentially in terms of spatial contribution and are incorporated into the plexus through mechanisms governed by different regulatory signaling pathways (Chen et al., 2014a; Payne et al., 2019; Sharma et al., 2017; Wu et al., 2012; Zhang & Zhou, 2013). Likewise, it has been shown that the SV contribution occurs through sprouting angiogenesis (Red-Horse et al., 2010; Tian et al., 2013a; Chen et al., 2014a; Sharma et al., 2017), while for endocardial contribution different mechanisms have been proposed, including angiogenesis (Payne et al., 2019; Tang et al., 2022; Wu et al., 2012), endocardial budding (Red-Horse et al., 2010; Sharma et al., 2017) or trapping during myocardial compaction (Tian et al., 2014).

Our three-dimensional characterization demonstrates that coronary morphogenesis is the result of two simultaneous angiogenic sprouting processes, in different regions of the developing heart. Although sprouting from the SV is the predominant mechanism of cardiac vascularization, the endocardium has increasingly emerged as a very reactive cardiac compartment. Multiple EndoECs of the IVS sprout both towards the forming septum and towards the surface of the ventral side of the heart. Additionally, we show that ventricular EndoECs have the potential to respond to the angiogenic cues that drive SV-derived plexus progression, resulting in a wave of endocardial angiogenic activation, and contributing to around 20% of the total coronary endothelium. While the endocardial contribution to ventral and IVS plexus is generally accepted (Chen et al., 2014a; D’amato et al., 2022; Phansalkar et al., 2021), the contribution to the FVWs plexus has been previously underestimated (Lu et al., 2021; Zhang et al., 2016).

Additionally, we have detected a bias of endocardial-derived cells to differentiate into arteries and capillaries, rather than veins. This could be explained by the preferential contribution of endocardial-derived vessels to the inner layer of the myocardial wall, where prospective arteries will form (Chen et al., 2014a; Wu et al., 2012). Consistently, we have detected ventricular EndoECs which show upregulation of CoEC markers and a transcriptional profile that partially overlaps with the profile of a particular subtype of CoECs (CoEC II), found at the inner myocardial layer with a capillary-to-artery signature.

Beyond providing firm proof for a substantial endocardial contribution to the intramyocardial plexus of the FVWs, our data demonstrate that this angiogenic event is responsible for the establishment of lumenized and perfused endocardial-coronary connections that might play a crucial role in plexus remodeling.

A recent study has identified endocardial tunnels that connect to the coronary vasculature at postnatal stages (Tang et al., 2022). These connections are detected shortly after birth, as extension of *Npr3-lineage* endocardium into the myocardial walls, which undergo an arterialization process upregulating Cx40 expression and recruiting mural cells. Finally, these endocardial-derived arteries establish connection with preexisting non-endocardial-derived coronary arteries. This novel mechanism has been proposed to be responsible for the drastic expansion of coronary arteries that occurs at postnatal stages and plays a protective role after ischemia.

Curiously, we detected many similarities between the postnatal establishment of these endocardial tunnels and the developmental endocardium-coronary connections we here describe. First, postnatal endocardial tunnels are initiated by an angiogenic process, that involves the upregulation of *Pdfgb* by EndoECs, as we have described during embryonic stages. Second, these connections undergo an arterialization process, transitioning from a more immature arterial state with no mural coverage and lack of Cx40 expression, into a more mature arterial state that recruits mural cells and upregulates Cx40. Similarly, we have detected, already during embryonic development, endocardial-coronary connections that undergo an arterialization process, involving the transition into the so-called ‘pre-artery’ state (Su et al., 2018). Whether all developmental endocardial connections are maintained, or whether they are all later remodeled into mature arteries, is a question that requires further investigation. Our data suggest that, at the IVS plexus, endocardial-coronary connections are resolved after first endocardial angiogenic phases, which has been described to occur before E13.5 (D’Amato et al., 2022). In contrast, the plexus of the FVWs is the result of a more prolonged angiogenic process that occurs up to E15.5. As a consequence, an increasing number of endocardial-coronary connections are detected up to this stage. Unfortunately, their high number and the large size of the samples, make it very difficult to quantify such connections in postnatal hearts. Although some pruning of endocardial-coronary connections is expected to occur in the FVWs, as it does in the IVS plexus, unequivocally they are still present at neonatal stages (fig. 6).

The establishment of structures similar to the endocardial tunnels had been previously described in adult hearts upon myocardial infarction (referred as ‘endocardial-flowers’), in a process that involves both endocardial angiogenesis and tunnel arterialization as well as the connection with preexisting coronary arteries (Miquerol et al., 2015). Similarly, perfused endocardial-coronary connections have been described in a model of VEGFB myocardial overexpression. In rats, VEGFB overexpression leads to the appearance of coronary arteries in the subendocardial zone, while no surrounding capillary proliferation was detected (Bry et al., 2010). In mouse, endocardial-coronary perfused connections upon VEGFB overexpression are also observed, an event that improves reperfusion and cardiac function upon myocardial infarction (Räsänen et al., 2021).

On the basis of these results, we propose that endocardial-coronary connections are a general and conserved mechanism, described here for the first time during embryonic development, but that also occurs during the postnatal expansion of the coronary vasculature (Tang et al., 2022) and in adult, upon ischemia (Miquerol et al., 2015) or endocardial activation regimes (Bry et al., 2010; Räsänen et al., 2021). Endocardial-coronary connections are initiated through angiogenic sprouts from the endocardium, which anastomose with the coronary plexus and undergo arterialization transitioning to a pre-artery state before they specify into mature arteries. Our single cell analysis identified putative endocardial progenitors of coronary EC, which are not exhausted during coronary development, and may retain their potential to engage in *de-novo* sprouting later in life. We propose that these cells will be the source of endocardial angiogenesis also in adult hearts upon reactivation situations.

The current major challenge is to elucidate the mechanisms underlying the arterialization process of the endocardial-coronary connections, but also of the coronary arteries of the myocardial wall, since both involve a pre-arterial specification step.

In the mouse retina as well as in the zebrafish embryo, pre-arterial specification has previously been proposed to be linked to tip cell sprouting. In these vascular beds, Dll4-Notch signaling interactions at the sprouting front induces an endothelial fate switch and specifies future arterial cells among the tip cell progeny (Fang et al., 2017; Hasan et al., 2017; Hou et al., 2022; Pitulescu et al., 2017; Xu et al., 2014). More recent evidence points to a priming mechanism of tip cells towards an arterial fate that is linked to the Notch pathway directly inducing metabolic and cell cycle arrest (Chavkin et al., 2022; Fang et al., 2017; Luo et al., 2021; Su et al., 2018) and subsequently more responsiveness to migratory cues (Hasan et al., 2017; Luo et al., 2021; Pitulescu et al., 2017). The Cxcl12-Cxcr4 chemokine signaling axis has been identified as a critical downstream target of Notch in this process (Hasan et al., 2017; Luo et al., 2021; Pitulescu et al., 2017). Based on our findings, we propose that pre-arterial specification is subordinate to angiogenic sprouting also in the heart. First, we have demonstrated that the inner myocardial wall is vascularized by sprouting from both SV-derived subepicardial vessels as well as from the endocardium. Second, we have identified that cardiac pre-arterial cells, beyond having a dual capillary and arterial transcriptional profile, also exhibit a strong tip cell marker signature. At the same time, they show diminished cell cycle activity, at similar levels as differentiated arterial cells, which has been recently described as a prerequisite to trigger endothelial cell arterialization (Luo et al., 2021). Additionally, our histological studies identified the appearance of Cxcr4-expressing pre-arterial cells in the inner myocardial wall as soon as this area is vascularized - by sprouting. As the plexus is remodeled, these Cxcr4-expressing cells become absent from the capillaries and are only found in the forming arteries, confirming that pre-arterial cells coalesce to eventually form arteries.

Further evidence linking pre-arterial specification and sprouting also in the heart stems from the display of analogous vasculature phenotypes in heart and retina when key regulatory players of sprouting are altered. First, in the retina, endothelial loss of Dll4 leads to hypersprouting that results in a dramatic increase of vascularization coupled to impaired arterial patterning (Hellström et al., 2007; Pitulescu et al., 2017; Suchting et al., 2007). In the heart, loss of Dll4 in CoEC leads to a denser ventricular coverage of the coronary plexus, more pronounced at the inner myocardial wall, and defective arterial specification (Lu et al., 2021; Travisano et al., 2019). Second, in the retina, Cxcr4 is expressed in tip cells and arterial cells, confirming the Cxcr4 pattern we see in the developing heart. Also, endothelial loss of Cxcr4 in the retina leads to reduced sprouting and vessel progression. In the heart, it also leads to defective intramyocardial vascularization and defective coronary artery formation, which results in a thin ventricular wall phenotype (Cavallero et al., 2015; Ivins et al., 2012). Furthermore, inducible endocardial loss of Cxcr4 is sufficient to impair coronary artery formation (D’amato et al., 2022).

Surprisingly, while coronary artery formation is impaired in these conditions, subepicardial plexus and vein development is normal, suggesting sprouting within the subepicardium as well as venous remodeling may be governed by different mechanisms than sprouting towards the myocardial wall.

Our results further indicate that perfusion of the coronary plexus and venous specification occur already prior to the connection to the aorta. Other studies in avians are in agreement with an early perfusion of the network before the aorta connection (Vrancken Peeters et al., 1997; Palmquist-Gomes et al., 2018). Further investigation is required in order to decipher whether this low shear-stress perfusion is enough to trigger venous remodeling. On the other hand, despite of the plexus being perfused and pre-arterial cells being specified before the formation of the coronary ostia, the final remodeling of the coronary arteries requires the onset of an unidirectional and high-shear flow from the aorta. This event triggers the increase of the intramyocardial vessel diameter (fig. S8), and the recruitment of mural cells (Volz et al., 2015). Indeed, many studies have demonstrated that impaired coronary ostia formation results in lack of arterial remodeling (Cavallero et al., 2005; Chen et al., 2014b; Gonzalez-Iriarte et al. 2003; Icardo et al. 2001; Ivins et al., 2015; Walker et al. 2005).

Taken together, we propose a model of coronary plexus formation and arterial remodeling (fig. 7), where multiple angiogenic events from different sources, first, give rise to an integrated coronary plexus, and second, determine the subsequent remodeling of the network, either by the specification of pre-arterial cells during sprouting or by the establishment of coronary-endocardial connections that are arterialized.

**Figure 7.**
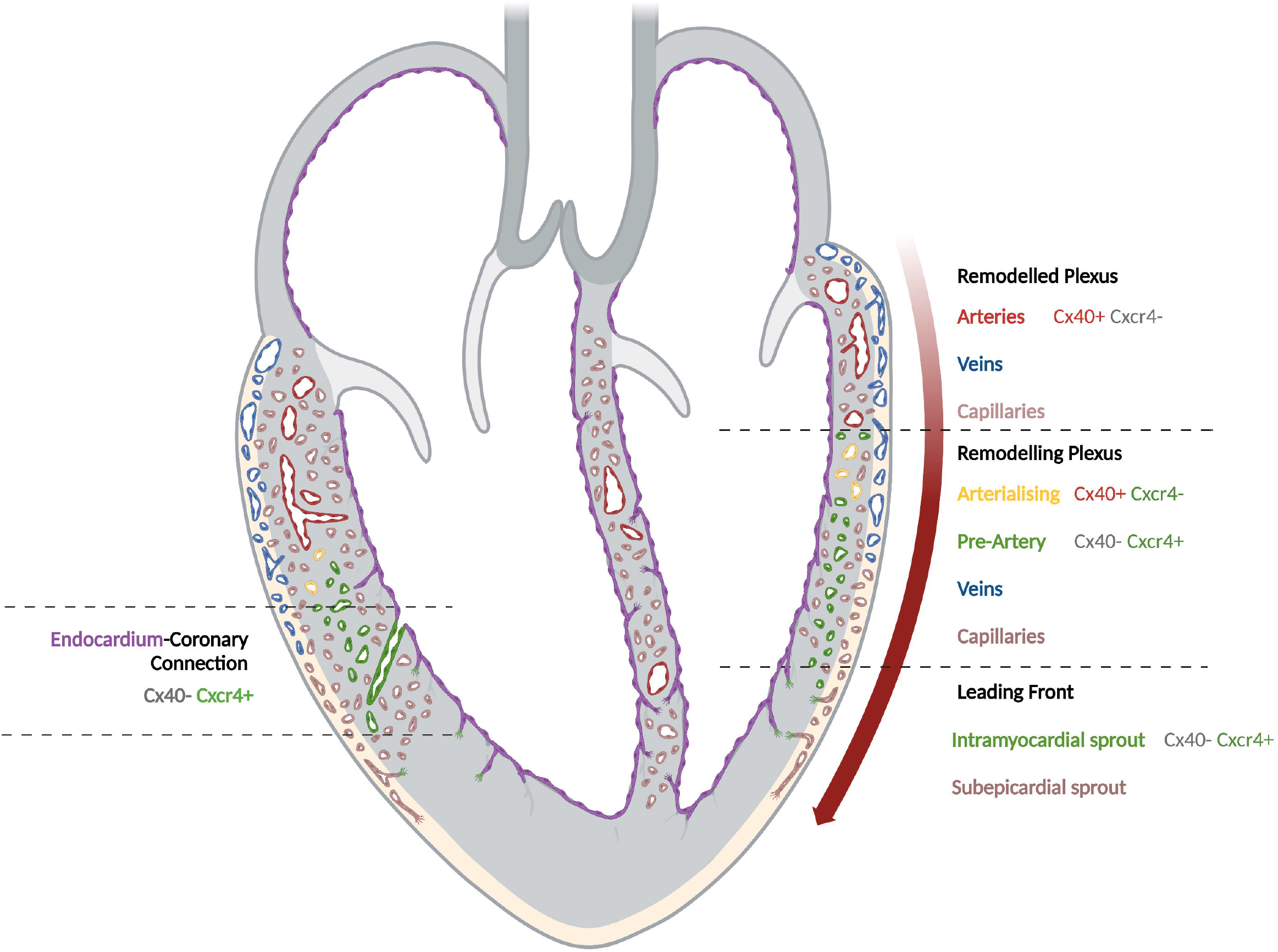
Abstract Figure. Schematic a longitudinal section of an embryonic heart. On the right side, the SV-derived coronary plexus (brown) expands within the subepicardium (beige) where veins will be formed (blue). As plexus progresses along the subepicardial space (Cxcr4-tip cell, brown), the intramyocardial wall (gray) is vascularized by sprouting (Cxcr4+ tip cells, green) of both subepicardial plexus (brown) and endocardium (purple). Sprouting tip cells towards the inner myocardium are specified as pre-arterial (Cxcr4+ cells, green). Upon flow onset (E13), they undergo arterialization (Cxcr4+Cx40+, yellow) and finally remodeled into mature arteries (Cx40+, red). On the left side, endocardial angiogenesis leads to endocardium-coronary vessels connections that show signs of arterialization (Cxcr4+, green).

Understanding the mechanisms underlying the vascularization of the heart, not only during angiogenic phases, but also during its specification into a more mature hierarchical vascular network, is key in order to promote therapeutic neovascularization strategies upon ischemic injury. Tailoring such strategies to distinct vascularization mechanisms for the intramyocardial and subepicardial vascular structures may soon become feasible as more insights into the distinct regulatory and morphogenic principles emerge.

## Methods

### Mouse strains and tamoxifen-induced lineage tracing

The following mouse strains were used: *PdgfbCreERT* (Tg(Pdgfb-icre/ERT2,-EGFP)1Frut) (Claxton et al., 2008), *BmxCreERT* (Tg(Bmx-cre/ERT2)1Rha) (Ehling et al., 2013) and *R26mTmG* (Gt(ROSA)26Sortm4(ACTB-tdTomato,-EGFP)Luo/J) (Muzumdar et al., 2007). Mice were maintained at the Max Delbrück Center for Molecular Medicine under standard husbandry conditions and were handled in compliance with the institutional and European Union guidelines for animal care and welfare.

All embryos were staged from the time point of vaginal plug, which was designated as E0. Single injections of tamoxifen (Sigma-Aldrich) were performed intraperitoneally (10 μg/g animal) 24 hours prior analyzed time-point for *PdgfbCreERT;R26mTmG* and at E9.5 for *BmxCreERT;R26mTmG*. Embryos/pups were then collected from E10.5 to P0. Animal procedures were performed in accordance with the animal license G0246/17.

### Immunofluorescence staining

Embryos and pups were fixed in 2% fresh paraformaldehyde (PFA) solution in PBS overnight rocking at 4°C, washed in PBS, cryoprotected in sucrose solutions, embedded in optimum cutting temperature (OCT) embedding compound (Tissue-Tek), and frozen at −80°C. Tissue sections were first incubated with blocking buffer (10% donkey serum, 0.1% Triton X-100 and 1% BSA in PBS) for 1 hour at room temperature and then with primary antibodies overnight at 4°C. The following primary antibodies were used: anti-Cxcr4 (Abcam, ab124824), anti-Endomucin (Santa Cruz, sc-65495), anti-ERα(Santa Cruz, sc-543), anti-GFP(Abcam, ab13970), anti-Unc5b (Cell Signaling, 13851), anti-VeCadherin (BD Biosciences, 555289). Alexa fluor-conjugated secondary antibodies (Invitrogen) were incubated for 1-2 hours at room temperature to detect the signals. For some weak signals, HRP-conjugated secondary antibodies and tyramide signal amplification kit (PerkinElmer) were used to magnify the signals. Sections were mount with Vectashield (Vector). All images were captured on a Zeiss LSM-780 confocal microscope. ImageJ software was used as image analysis tool.

### Three-dimensional imaging

Clearing of the dissected embryonic or postnatal hearts were performed with an adapted iDISCO protocol (Renier et al., 2014).

Embryonic or postnatal hearts were dissected from the body and fixed in fresh 2% PFA overnight at 4°C and then thoroughly washed with PBS. Permeabilization was performed with PBT 0.2% for 2h, and then with PBS/0.1% Tween20/0.1% TritonX-100/0.1% Deoxycholate/0.1% NP40/20% DMSO overnight at room temperature. Samples were then blocked with blocking buffer (PBS/0.2%TritonX-100/10%DMSO/0.3Mglycine/6% Donkey) Serum for 12h at room temperature and washed with PBTwH(PBS/0.2% Tween-20 with 10ug/ml heparin) 1h. The incubation of primary antibody was performed in PBTwH / 5% DMSO/ 3% Donkey Serum, shaking for 24h at room temperature. The antibodies used were the same antibodies used for immunofluorescence on sections. The concentration of the primary antibody was 1:25-1:200 depending on the sample size. Afterwards, hearts were washed with PBTwH 1h shaking at room temperature eight times. Alexa Fluor-conjugated secondary antibodies were incubated in PBTwH/3% Donkey Serum overnight shaking at room temperature, at a concentration of 1:50-1:200 depending on the sample size. After thoroughly washing samples with PBTwH again 1h shaking at room temperature eight times. For clearing, samples were dehydrated with increasing concentrations of Tetrahydrofuran solution in H20 (THF): 1h in THF 50%, 1h THF 80% and two times 1h THF 100%, and then in Dicholoromethane until samples sink at the bottom (with a maximum of 5 minutes). Finally, samples were cleared in DiBenzyl Ether (DBE) shaking until the sample was clear. Samples can be stored in DBE at 4°C until imaged.

For imaging, samples were placed in a drop of DBE on a glass-bottomed dish. Image acquisition was performed with a long working-distance oil-immersion 25x objective on a Zeiss LSM-780 microscope. Z-stacks of a maximum of 460um were acquired. For samples bigger than E12, two different z-stacks acquisition were performed, one from de dorsal side of the heart, and the other from the ventral side.

### Three-dimensional images post-processing

After confocal images acquisition, Zen 3.4 Blue edition software (Zeiss) was used for stitching. Arivis Vision4D software (Zeiss) was used for registering dorsal and ventral *Z*-stacks when needed. ImageJ was used for manual segmentation. Imaris 9.9.0 (Oxford Instrumentals) was used for three-dimensional rendering, colocalization analysis and area quantification.

### Live imaging on explanted hearts

E12.5 *PdgfbCreERT;R26mTmG* embryos were dissected in PBS supplemented with 10%FBs, penicillin (100 U/ml) and streptomycin (100ug/ml). Hearts were isolated and transferred to Hepes-buffered DMEMF12 medium supplemented with 2% FBS and penicillin-streptomycin. Hearts then were embedded into a 0.5% low melting point agarose gel (Sigma-Aldrich) in a glass-bottomed culture dish (MatTek) and bathed in the same medium used for heart dissection. Images were captured in a 3i spinning-disc confocal microscope every 10 min for 15 hours. During image capture the culture chamber was maintained at 37 °C in a 5% CO2 humidified atmosphere.

### Single cell suspension preparation and cell sorting

Single-cell suspension of endothelial cells was prepared as described previously (Pinto et al., 201). C57Bl6 8 weeks old adult animals, P2 postnatal, E15 and E12 were used for single-cell suspension preparation. Adult animals were sacrificed by cervical dislocation while embryonic and postnatal animals were decapitated. Adult and postnatal hearts were chopped with a scalpel until getting 1-2mm pieces of tissue. The enzymatic digestion was adapted to every stage analyzed: for E12, 20min of Collagenase/Dispase (Sigma-Aldrich) (1mg/ml), for E15, 30min of Collagenase/Dispase (1mg/ml), and for P2 and adult hearts, 20 min of Collagenase II (500units/ml) (Worthington) plus a second digestion step with Collagenase/Dispase for 30min. All steps were performed at 37°C, under agitation and pippeting up and down the solution within the tubes every 5min. To stop the enzymatic reaction 10ml of cold FAC buffer (HBSS/2%FBS/10mM Hepes) was added to the samples and then spinned down (400G, 5 min). For adult samples, red blood cells lysis step (Invitrogen) was performed. After enzymatic digestion, cell debris removal kit (Miltenyi) was performed and finally, all suspension filtered through a 40um strainer.

Single-cell suspensions were incubated with primary antibodies in FACS buffer for 20 min on ice. The following antibodies were used: APC-conjugated anti-CD31 (Thermo Fisher, 17-0311-80) and PE-conjugated anti-CD45 (BD Biosciences, 553081). Samples were centrifuged (400G, 5 min) and resuspended on FACS buffer and stored on ice until applied to the flow cytometer. CD31+/CD45-cells were sorted on an Aria II into 1.5ml tubes with EBM2 media. Compensation controls were set up for each single channel. After completion of sorting, 10% DMSO was added and cell suspension frozen at −80°C. Several rounds of single-cell suspension preparation, sorting and freezing was repeated for embryonic samples in order to get a significant number of cells: 8 weeks old adult animals (n=6), P2 postnatal (n=21), E15 (n=48) and E12 (n=37).

Frozen vials of sorted CD31+/CD45-cells from E12, E15, P2 and 8wo adult hearts were thawed in a water bath at 37°C. For E12 and E15, a dead cell removal kit (Miltenyi) was applied. For P2 and adult, re-sorting by flow cytometry and selection of live cells was performed.

### Library preparation and 10X Chromium sequencing

Cell suspension was adjusted to 400-1000 cells per microlitre and loaded on the Chromium controller (10X Genomics) with a targeted cell recovery of 8000-10000 per reaction. 3’ gene expression libraries were prepared according to the manufacturer’s instructions of the v2 Chromium Single Cell Reagent Kits (10X Genomics). Quality control of cDNA and final libraries was done using Bioanalyzer 2100 (Agilent). Libraries were sequenced using HiSeq 4000 (Illumina) targeting a depth of 25000 reads per cell.

### Transcriptome mapping

After sequencing, the sequence data were mapped to the mouse reference genome (mm10 pre-build references v 2.1.0) provided by 10X Genomics using the CellRanger suite (v.2.2). The count matrices generated by CellRanger as followings were used for further analysis. Mapping quality was assessed using the CellRanger summary statistics.

### Empty droplets removal and doublet estimation

Empty droplets were identified by Emptydrops, which is implemented in the CellRanger workflow. After removal of empty droplets, we applied scrublet per sample (ref: 10.1016/j.cels.2018.11.005) to assign a doublet score (scrublet_score) to the metadata container of each cell.

### Cell quality control and filtering

Downstream analysis employed the concatenated filtered feature-barcode matrices, using Seurat (vref: 10.1016/j.cell.2021.04.048). Genes were filtered out when they were expressed in less than three cells. The filtering cut-off was decided based on the distribution of data quality and the estimated doublet ratio provided by 10X Genomics. We applied individual cell filtering criterion: cells from E12 were filtered for counts (nCount_RNA <= 4000), genes (nFeature_RNA <= 1500), mitochondrial genes (percent.mt <= 10%), and scrublet score (scrublet_score <= 0.15). Cells from E15 were filtered for counts (nCount_RNA <= 3000), genes (nFeature_RNA <= 1300), mitochondrial genes (percent.mt <= 10%), and scrublet score (scrublet_score <= 0.15). Cells from P2 were filtered for counts (nCount_RNA <= 12000), genes (nFeature_RNA <= 4000), mitochondrial genes (percent.mt <= 20%), and scrublet score (scrublet_score <= 0.15). Cells from Adult were filtered for counts (nCount_RNA <= 5000), genes (nFeature_RNA <= 2000), mitochondrial genes (percent.mt <= 20%), and scrublet score (scrublet_score <= 0.15).

### Cell cycle score calculation

Cell cycle score were calculated using CellCycleScoring function integrated in Seurat. Previously reported cell cycle genes (ref 10.1126/science.aad0501) are used for the reference of cell cycle genes after converting the human gene list into mouse orthologues using biomaRt (v2.52.0).

### Dimensionality reduction, clustering, and analysis of differentially expressed genes

After read count normalization and log2-transformation, top 2000 highly variable genes were selected. Then, regression of UMI counts, percent mitochondrial gene expression and cell cycle gene expression was performed. Prior to manifold construction using UMAP, top 30 principal components were harmonized by anchored canonical correlation analysis (CCA) (ref 10.1016/j.cell.2019.05.031). Cells were clustered using the original Louvain algorithms (ref 10.1140/epjb/e2013-40829-0). For all datasets, non-endothelial subtypes (e.g. blood and immune cells, cardiomyocytes, smooth muscle, fibroblasts) as well as a small number of lymphatic cells were removed. Likewise, immediate early genes-expressing cells were also removed as considered stressed cells during the single-cell suspension preparation process.

Differential expressed genes (DEGs) per cluster were calculated using FindAllMarkers function of Seurat with the Wilcoxon rank sum test. For the marker gene computation, we selected genes expressed in at least 25% of cells in either of the populations and with a log2-transformed fold change of at least 0.25. Genes with adjusted p-value < 0.05 were called as DEGs.

## Acknowledgments

We thank all members of the Gerhardt lab for interesting discussions and comments, and specially to Alexandra Klaus-Bergmann and Katja Meier for further technical support. We also thank Marie Altmann and the rest of the Mouse Facility staff at the MDC for excellent animal care, as well as to the Advanced Light Microscopy & Image Analysis technology platform at MDC for helping stablishing the imaging pipeline. We gratefully acknowledge Ralf Adams for sharing *BmxCreERT* mouse strain.

## Abbreviations

AVC: Atrioventricular Canal
CoECs: Coronary Endothelial Cells
CreERT: Cre recombinase – Estrogen Receptor fusion protein
E: Embryonic day
ECs: Endothelial Cells
Endoc: Endocardium
EndoECs: Endocardial Cells
FACS: Fluorescence-Activated Cell Sorting
FVWs: Free Ventricular Walls
GO-Term: Gene Ontology Term
IVS: Interventricular Septum
LV: Left Ventricle
P: Postnatal day
Prolif: Proliferating cells
OFT: Outflow Tract
RV: Right Ventricle
scRNAseq: Single-cell RNA sequencing
SV: Sinus Venosus
tdTom: tdTomato Fluorescent Protein
UMAP: Uniform Manifold Approximation and Projection

---

**Figure S1. Methods summary**

A. Lineage-tracing strategy used for three-dimensional imaging and scRNAseq.

B. Three-dimensional imaging pipeline.

RA=Right Atrium; LA=Left Atrium; RV= Right Ventricle; LV=Left Ventricle, AVC=atrioventricular canal, IVS=interventricular septum, SV=sinus venous (inflow tract), OFT=outflow tract.

**Figure S2. *Pdgfb-iCreERT* is a suitable lineage-tracing tool for coronary endothelium**

A. Percentage of coronary endothelium that express *Pdgfb* in the atrial plexus along development. Quantified as GFP/Endom+VeCadh colocalization area (μm^2^), normalized to Endom+VeCadh total area, on the segmented coronary plexus of both atria (n=3).

B. Percentage of coronary endothelium that express *Pdgfb* (normalized GFP/Endom+VeCadh colocalization area) in the ventricular plexus along development (n=3).

C. Percentage of GFP-expressing cells that also express Fabp4 (left) and percentage of Fabp4-expressing cells that also express GFP (right) in the whole mount heart. Quantified as GFP/Fabp4 colocalization area (μm^2^), normalized to either GFP (left) or Fabp4 (right) total area (n=3).

D. Section of a E15.5 *PdgfbCreERT;R26mTmG heart* stained against Fabp4 (blue) and Endom+VeCadh (magenta).

E. Percentage of endocardium that express *Pdgfb* (normalized GFP/Endom+VeCadh colocalization area) in the valves along development (n=3). Section showing a representative valve of a E13.5 *PdgfbCreERT;R26mTmG heart*.

F. Percentage of OFT endothelium that express *Pdgfb* (normalized GFP/Endom+VeCadh colocalization area) along development (n=3). Section showing a representative OFT of a E13.5 *PdgfbCreERT;R26mTmG heart*.

G. Three-dimensional rendering of the IVS of a E11.5 *PdgfbCreERT;R26mTmG heart*,showing early angiogenic response of the endocardial zone of this region.

H. Three-dimensional rendering of the IVS of a E11.5 *PdgfbCreERT;R26mTmG heart*, showing large angiogenic activity. All endocardial protrusions are lumenized, some protrude directly into the subepicardium and often form buds (arrowheads). F” is same three-dimensional image as F’ rotated vertically 180°.

I. Three-dimensional rendering of the ventral side of the LV of a E15.5 *PdgfbCreERT;R26mTmG heart*, showing anastomosing plexuses. Before the subepicardial area of the LV ventricle is not vascularized, the inner myocardial plexus is formed.

RA=Right Atrium; LA=Left Atrium; RV=Right Ventricle; LV=Left Ventricle. SV=sinus venous, Subepic=Subepicardium, Intramyo=Intramyocardium. Scale bars=100 μm in C, I; 25μm in E, F, G, H; 10μm in F’, F”. Data are mean ± SD, ns=not significant, *=p<0.05, **=p<0.01, ***=p<0.001, by Welch’s t-test.

**Figure S3. *Pdgfb*CreERT recombination window only occurs during 48h upon tamoxifen administration.**

A-E. Sections of E11.5-E15.5 *BmxCreERT;R26mTmG* hearts, stained against ERα to analyze the distribution of the CreERT protein (under *Bmx* promoter) along development. Black arrowheads point to GFP-negative/tdTom-positive cells; yellow arrowheads to GFP-positive/tdTom-positive cells; white arrowheads to GFP-positive/tdTom-negative/ERα-positive cells; and cyan arrowheads to GFP-positive/tdTom-negative/ERα-negative cells.

F. Schematic representation of GFP, tdTom and ERαsignals detected in panels above. By analyzing the colocalization of GFP and tdTom we observe recombination exclusively in endocardial cells and only during the first 48h upon tamoxifen administration. Also, in this time window, ERαlocalizes into the nuclei of *Bmx*-expressing endocardial cells. From E12.5 onwards, ERαis found in the cytoplasm of all *Bmx*-expressing endocardial cells. In capillaries, we never observe CreERT or tdTom expression, indicative that GFP labeling is because they are derived from the *Bmx*-lineage endocardium.

In arteries, we detect ERα expression, which recapitulates *Bmx* expression. However, ERα is always located at the cytoplasm, so the GFP labeling in arterial ECs is not due to autonomous recombination. Also, we observe ERα-expressing arterial cells that lack GFP, evidencing the tamoxifen is not leading to any recombination event at this stage. Scale bars=100μm; 10μm in boxed areas.

**Figure S4. *Pdgfb*CreERT recombination window only occurs during 48h upon tamoxifen administration.**

A. Lineage-tracing strategy. Comparison of E15.5 *BmxCreERT;R26mTmG* hearts when tamoxifen is administered at different timepoints (E9.5 and E13.5).

B. Percentage of *Bmx*-lineage cells within endocardium or coronary plexus, segmented in capillaries, arteries or veins at E15.5. Quantified as GFP/Endom+VeCadh colocalization area (μm^2^), normalized to Endom+VeCadh total area. E13.5 tamoxifen administration leads to a decrease labeling of *Bmx*-lineage derived coronary plexus. (n=3-7, see dots on bars).

C. Section of a E15.5 *BmxCreERT;R26mTmG* heart, stained against αSMA and Endom+VeCadh. Only when tamoxifen is administered at E13.5, *Bmx*-lineage non-endothelial cells are observed (green outline).

Endoc=Endocardium, Tam=Tamoxifen. Scale bar=50μm. Data are mean ± SD, ns=not significant, *=p<0.05, ***=p<0.001, by Welch’s t-test.

**Figure S5. Single-cell sequencing identifies putative endocardial progenitors of coronary endothelium**

A. Gene expression profile of *Car4* and *Cxcl12* in the UMAP plot.

B-C. Different point of views of the three-dimensional projection of UMAP shown in Fig. 5A (EC clusters from E12-E15-P2-Adult combined datasets)

C. GO-term analysis of Endoc I and Endoc IV.

D. Differential expressed gene analysis of Endoc I and Endoc IV versus all other endocardial clusters (Endoc II, Endoc III and Endoc V).

E. Three-dimensional rendering of a E14.5 *BmxCreERT;R26mTmG* heart stained against Fabp4 (blue) and Endom and VeCadh (magenta). Please note that Fabp4-expressing cells are also expressing GFP (green). Scale bars=100μm.

F. Gene expression profile of *Gja5* (Cx40) in the UMAP plot split by every timepoint studied.

**Figure S6. Validation of annotated clusters by *in situ hybridization* of top expressed gene markers**

A. For each cluster, its distribution on the UMAP plot and representative ISH images of three top expressed genes from E14.5 wildtype embryos are shown. For each gene, together with the ISH images, it is shown the pattern of expression in the UMAP plot and a summary schematic figure of the observed ISH signal. ISH images are from GenePaint database (Visel et al. 2014).

**Figure S7. Pre-arterial cells immunolocalization at developmental stages**

A. Left-to-right serial sections of a E15.5 *BmxCreERT;R26mTmG* heart, stained against Unc5b (red) and Endom+VeCadh (blue). In the bottom panel, Unc5b expression is showed in gray scale for better visualization.

RA=Right Atrium; LA=Left Atrium; RV=Right Ventricle; LV=Left Ventricle, IVS=Interventricular septum. PA=Pulmonary artery, Ao=Aorta. Scale bars=100 μm.

**Figure S8. Coronary plexus is perfused and remodeled before the connection to the aorta**

A-C. Three-dimensional rendering of E11.5 (A), E12.5 (B) and E13.5) (C) *BmxCreERT;R26mTmG* hearts stained against ICAM2 (magenta) and corresponding single z planes. A’, B’ and C’ show a maximum intensity projection of the aorta in order to visualize whether coronary plexus is connected. Since early formation of coronary plexus, vessels are already lumenized and perfused. Red blood cells (asterisks) are observed in the lumen of coronary vessels and endocardial buds (gray arrowheads). Despite first signs of arterial remodeling (white arrowheads) is coupled to the remodeling of the upwards coronary stems (cyan arrowheads, compare A’and B’with C’), venous remodeling (black arrowheads) is earlier detected.

RA=Right Atrium; LA=Left Atrium; RV=Right Ventricle; LV=Left Ventricle, Ao=Aorta, ACV=Anterior coronary vein, CA=Coronary artery. Scale bars=100 μm; 10μm in boxed areas, A’,B’C’.

